# PGC-1α in the myofibers regulates the balance between myogenic and adipogenic progenitors affecting muscle regeneration

**DOI:** 10.1101/2021.11.03.466970

**Authors:** Marc Beltrà, Fabrizio Pin, Domiziana Costamagna, Robin Duelen, Alessandra Renzini, Riccardo Ballarò, Lorena Garcia-Castillo, Ambra Iannuzzi, Viviana Moresi, Dario Coletti, Maurilio Sampaolesi, Fabio Penna, Paola Costelli

## Abstract

Skeletal muscle repair is accomplished by satellite cells (MuSC) in cooperation with interstitial stromal cells (ISCs). So far, the relationship between the function of these cells and the metabolic state of myofibers remains unclear. The present study reports alterations in the proportion of both MuSCs and adipogenesis regulators (Aregs) induced by overexpression of peroxisome proliferator-activated receptor gamma coactivator 1-alpha (PGC-1α) in the myofibers (MCK-PGC-1α mice). Although PGC-1α–driven increase of MuSCs does not accelerate muscle regeneration, myogenic progenitors isolated from MCK-PGC-1α mice and transplanted into intact and regenerating muscles are more prone to fuse with recipient myofibers than those derived from WT donors. Moreover, both young and aged MCK-PGC-1α animals show reduced perilipin-positive areas when challenged with an adipogenic stimulus, demonstrating low propensity to accumulate adipocytes within the muscle. These results provide new insights on the role played by PGC-1α in promoting myogenesis and hindering adipogenesis in the skeletal muscle.

## INTRODUCTION

Muscle regeneration is a complex multi-step process that relies on the satellite cells (MuSCs) responsible for the postnatal myogenesis, as well as the maintenance of muscle integrity^1^. Under normal circumstances, MuSCs are quiescent and quickly activate upon injury, dividing and differentiating into myoblasts, that ultimately fuse to generate new myofibers. Although muscle regeneration is dependent upon MuSCs, it also requires the participation of other non-myogenic cells involved in orchestrating inflammation, debris clearance, extracellular matrix deposition and extrinsic regulation of MuSC activity^2^. In particular, interstitial stromal cells (ISCs) can participate to adult myogenesis and support MuSC function. The remarkable regenerative capacity of the skeletal muscle is compromised in aging due to decreased number and function of MuSCs, contributing to sarcopenia and frailty^3–5^. Aged-related faulty muscle regeneration is partially attributed to the reduced ability of ISCs, in particular of fibro-adipogenic progenitors (FAPs), to support MuSC activation and differentiation^6^. Moreover, functional decline of accessory cell is characteristic of myopathies such as Duchenne muscular dystrophy^7^ and systemic conditions such as obesity, diabetes and cancer cachexia^8–11^.

Mitochondria are critical for preserving the metabolic fitness of the skeletal muscle. These organelles are deeply interconnected and form a network regulated by dynamic processes involving mitochondrial biogenesis, fission, fusion and mitophagy^12^. Upon activation and differentiation, MuSCs undergo a major genetic reprogramming to become metabolically active. Shortly after exiting quiescence, mitochondrial genes are robustly induced and MuSCs quickly accumulate mitochondrial mass in order to support the increasing energy demand needed for cell proliferation and myotube formation^13,14^. Mitochondrial dysfunction is a distinctive aspect of MuSC senescence in aging^15^. Consistently, preventing the decline of mitochondrial metabolism during aging rescues MuSC myogenic potential^15,16^, demonstrating that mitochondria are essential for the functional maintenance of MuSCs. Among the known players contributing to mitochondrial homeostasis, the peroxisome proliferator-activated receptor gamma coactivator 1α (PGC-1α) stands out as powerful driver of mitochondrial biogenesis^17^ and regulator of processes such as mitochondria fusion-fission events and mitophagy^18–22^. Aside from promoting muscle oxidative metabolism, PGC-1α induces angiogenesis^23^, neuro-muscular junction remodeling^24^ and increases the expression of structural proteins such as myosin heavy chain (MyHC) and utrophin^25^. Consistent with the broad effects on tissue plasticity, forced PGC-1α overexpression in several experimental models of atrophy preserves skeletal muscle function and myofiber morphology^26–32^.

In addition to its beneficial effects in the skeletal muscle, PGC-1α promotes the secretion of exercise-related myokines with both paracrine and endocrine functions, contributing to the crosstalk among muscle and fat, bone or brain^33^. Consistently, PGC-1α overexpression in mature myofibers impacts on the MuSC niche^34^, modulating the local pro- and anti-inflammatory cytokine balance^35,36^. Nonetheless, to our knowledge, no clear association exists between the predominant oxidative environment, as dictated by myofiber PGC-1α overexpression, and MuSC and ISC function. The present study provides new cues on the indirect impact of PGC-1α in altering the balance and propensity to differentiate of myogenic and adipogenic populations that reside in the skeletal muscle.

## RESULTS

### PGC-1α expression is transiently induced in early phases of muscle regeneration

Muscle regeneration is a highly energy-demanding process, and a progressive increase of mitochondrial function occurs in order to respond to the increased energetic needs^37^. Along this line, we characterized the reprograming of oxidative metabolism in the *tibialis anterior* (TA) muscle of wild-type (WT) mice after BaCl_2_-induced injury. Muscle regeneration was assessed at 7, 10 and 13 days post-injury (dpi), when new myofiber formation and maturation occurs, and at 49 dpi (7 weeks post-injury) as a final-phase myofiber maturation timepoint. Myofiber cross-sectional morphology was evidently disrupted at 7, 10 and 13 dpi, being associated with the presence of cell infiltrate, and was restored at 49 dpi, although central myonuclei still persisted (Figure 1A). Metabolic phenotype analysis, assessed by succinate dehydrogenase (SDH) staining, revealed a lack of definite phenotype at 7 and 10 dpi, with apparent improvement at 13 dpi and complete recovery at 49 dpi (Figure 1B). This trend was consistent with the initial loss and the subsequent progressive restoration of mitochondrial SDHA protein levels as muscle regeneration proceeds (Figure 1C, 1D). Interestingly, PGC-1α protein content was strongly increased at the initial stages of muscle regeneration (7 and 10 dpi), returning to normal levels at 13 and 49 dpi (Figure 1C, 1E). Consistently, expression of the PGC-1α downstream target gene *Cox4* was enhanced at 7 and 10 dpi (Figure 1F), although no changes in *Cycs* or *Sdha* expression was observed (Figure 1G, H). Notably, accumulation of PGC-1α protein levels correlated with the induction of myogenic markers *Pax7*, *Myog* and *Myh3*, and M1 macrophage marker *Cd68* at 7 and 10 dpi (Figure S1A-S1D). Regarding mitochondrial clearance, the mitophagy regulator gene *Prkn* was upregulated at 10 dpi (Figure 1I), whereas levels of *Fis1* and *Mfn2*, genes involved in mitochondrial fission and fusion respectably, remained unchanged (Figure 1J, 1K). Altogether, PGC-1α expression is induced as a consequence of myofiber formation and presumably promotes the expression of genes related to mitochondrial homeostasis during muscle regeneration.

**Figure 1.**
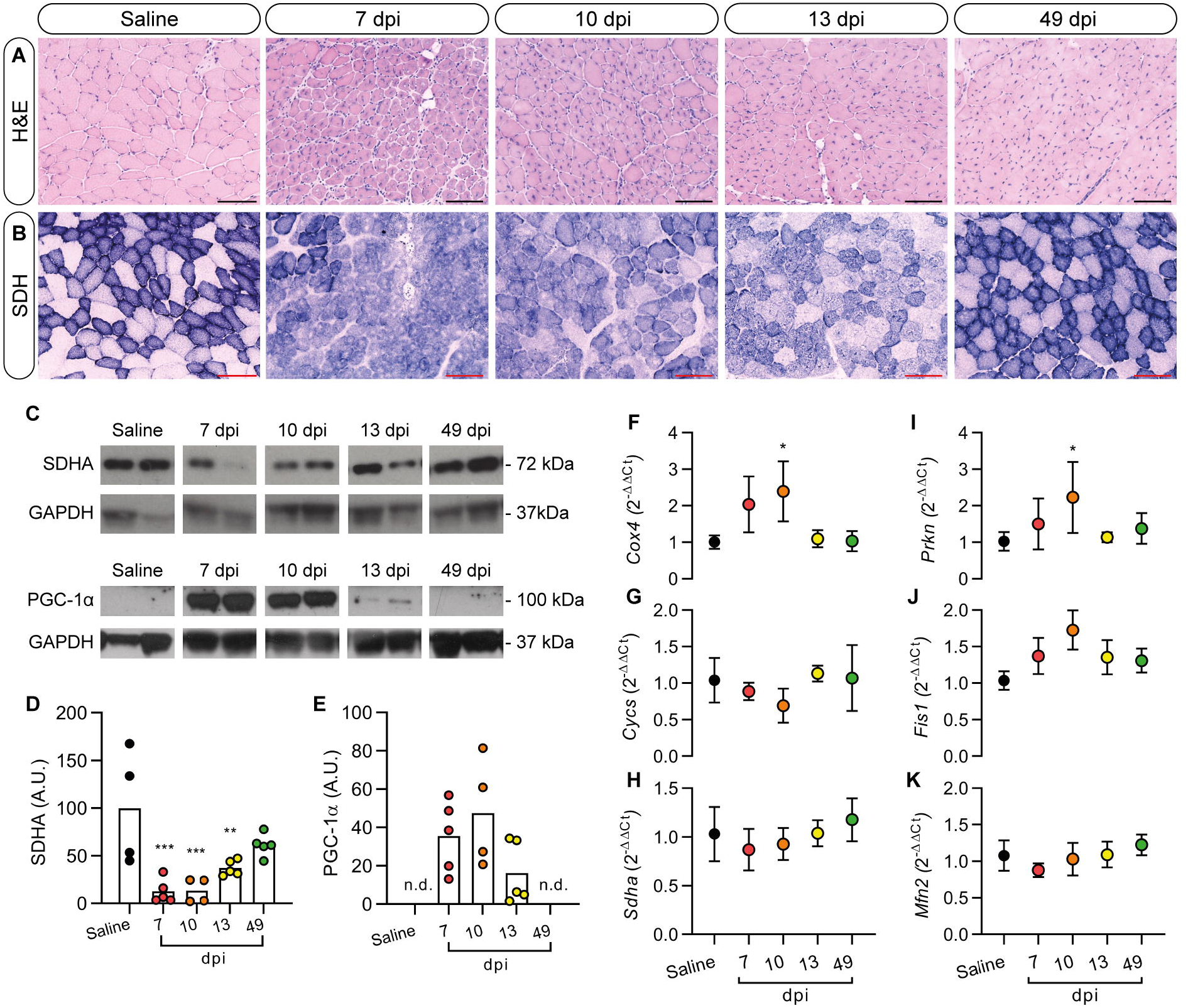
Modulation of mitochondrial markers during muscle regeneration. The TA muscle of WT mice was injured with 1.2% BaCl_2_ and muscle regeneration was assessed at 7, 10, 13 and 49 dpi. Every time point was assessed in n=5 mice except from 10 dpi with n=4 mice. Saline corresponds to the non-injured contralateral TA muscle of 49 dpi group (n=4); (A-B) Representative images of H&E and SDH staining of intact (saline) and BaCl_2_-injected TA muscles (scale bar: 100 μm); (C-E) Densitometry analysis of western blotting bands of PGC-1α and SDHA proteins. Total protein load was normalized by means of GAPDH protein expression; (F-K) RT-qPCR quantification of *Cycs*, *Cox4*, *Sdha*, *Prkn*, *Fis1* and *Mfn2* genes. Data are normalized by *Actb* expression and displayed as relative expression (2^−ΔΔCt^; mean ± SD) *vs* Saline group. Statistical analysis: *p<0.05, **p<0.01, ***p<0.001 *vs* Saline group (either One-way ANOVA + Dunnett’s test or Kruskal–Wallis + Dunn’s test).

### PGC-1α-driven metabolic switch enhances myogenic potential of progenitor cell populations

The marked increase of PGC-1α in regenerating muscles paved the way to investigate whether modulating this transcriptional cofactor could impinge on myogenesis. To this aim, MCK-PGC-1α mice were used, which constitutively overexpress PGC-1α in adult skeletal muscle (Figure S2A). Beyond the elevated mitochondrial content (Figure S2B, S2C), MCK-PGC-1α animals show an increased number of central myonuclei compared to WT animals in the absence of injury (Figure 2A), suggesting a constitutive activation of MuSCs. Consistent with this observation, enzymatic digestion of hindlimb muscles revealed a number of mononucleated cells in MCK-PGC-1α mice which was increased more than twice in comparison to WT animals (Figure 2B). In order to clarify if muscle-specific PGC-1α overexpression affected MuSC myogenic potential, myoblasts were isolated from two muscles with different metabolic phenotypes (the mostly glycolytic *extensor digitorum longus* (EDL), and the predominantly oxidative *soleus*) and cultured in differentiation medium. Myotubes from WT-derived EDL explant accumulated lower levels of MyHC compared to those obtained from WT-*soleus* (Figure 2C), demonstrating that muscle metabolic background likely impacts on the myogenic potential of MuSCs. Following this line, myotubes differentiated from MuSC isolated from the EDL muscle of transgenic mice showed increased MyHC levels in comparison to those derived from WT-EDL. By contrast, myotubes obtained from *soleus*-derived MuSC accumulated similar levels of MyHC independently from mouse genotype (Figure 2C).

**Figure 2.**
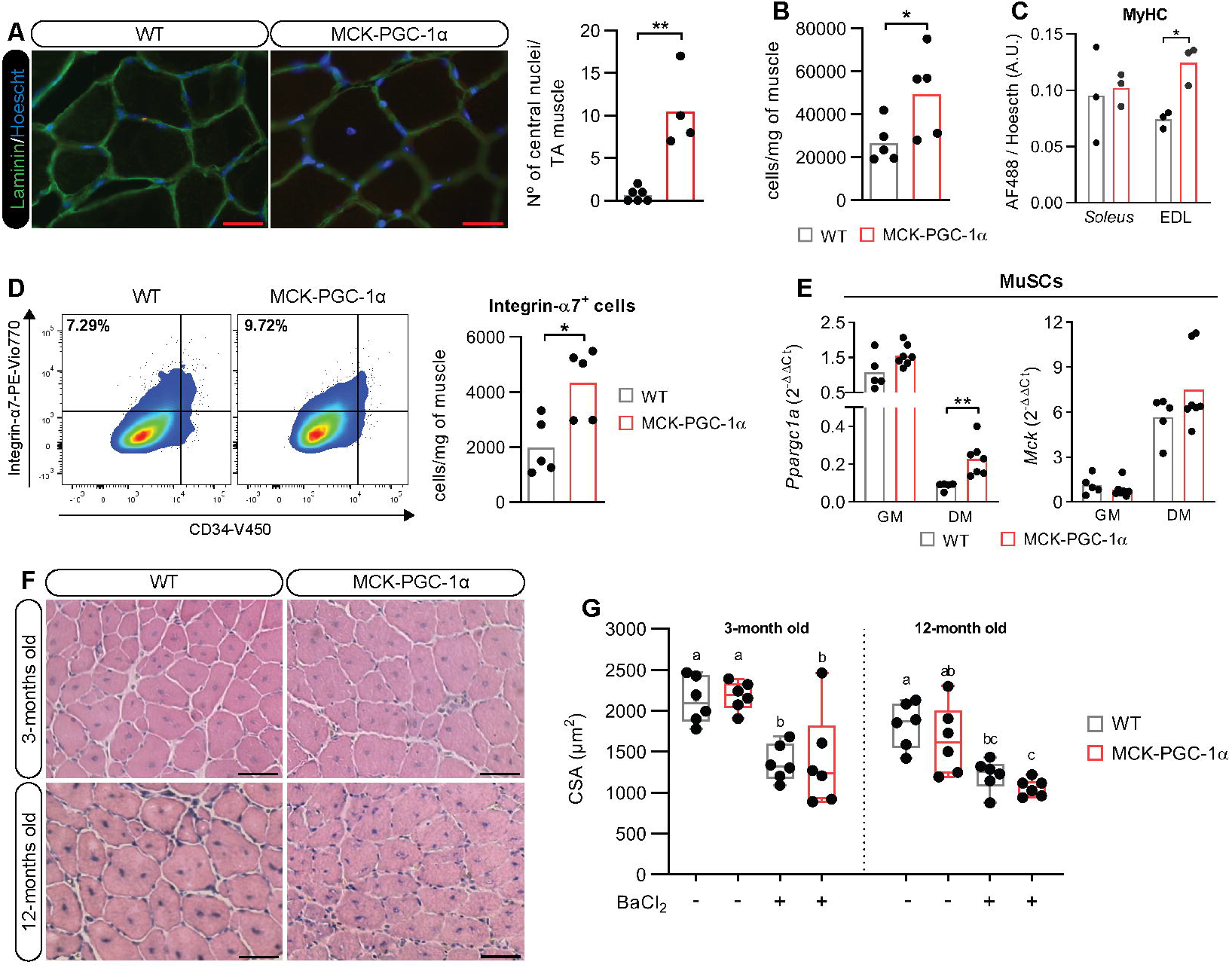
MCK-PGC-1α mice show increased myogenic precursors in the skeletal muscle, endowed with enhanced differentiation capacity *ex vivo* but not *in vivo*. (A) Representative images of Laminin/Hoechst immunostaining (scale bar: 25 μm) and quantification of central myonuclei of the TA muscle of WT (n=6) or MCK-PGC-1α (n=4) mice; (B) Total cell count after isolation of muscle interstitial cells from hindlimb muscles of WT (n=5) and MCK-PGC-1α (n=5) mice normalized per mg of tissue; (C) Quantification of AF488 fluorescent signal according to anti-MyHC antibody binding in the supernatant of lysates of *soleus* and EDL-derived primary myotube cultures of both WT (n=3) and MCK-PGC-1α mice (n=3); (D) Representative panels of gate distribution and relative quantification of positive events (cells) according to anti-integrin-α7 and anti-CD34 antibody binding in hindlimb muscles of WT (n=5) and MCK-PGC-1α (n=5) mice. Numbers in gates represent the mean percentage of cells by each specific labeling combination. Total number of MuSCs cells was obtained by normalizing integrin-α7 relative positivity with number of isolated cells/mg of tissue; (E) *Ppargc1a* and *Mck* gene expression in proliferating (GM) and differentiated (DM) MuSCs derived from WT (n=5) and MCK-PGC-1α (n=7) cultured *ex vivo*. Data are normalized by *H2bc4* expression and displayed as **relative expression** (2^−ΔΔCt^; mean) *vs* WT group; (F) Representative images of H&E staining (scale bar: 50 μm) of WT (n=6) and MCK-PGC-1α (n=6) animals injured using 1.2% BaCl_2_. Skeletal muscles were evaluated at 14 dpi; (G) CSA of TA muscles expressed as percentages of WT intact muscles. Significance of the differences: *p<0.05, **p<0.01 *vs* WT group (either Student’s t-test or Mann–Whitney test); groups with distinct letters are statistically different (Two-way ANOVA + Tukey’s test)

To better describe how PGC-1α overexpression impacts on myogenic populations, muscle interstitial cells were isolated, stained for integrin-α7 (a MuSC specific marker^38^) and characterized by flow cytometry. The analysis revealed that the skeletal muscle of MCK-PGC-1α mice contains more integrin-α7^+^ cells in comparison to WT ones (Figure 2D). The possibility that an artefactual overexpression of PGC-1α in MuSCs could occur was assessed. The results show that *Ppargc1a* expression was comparable in WT and MCK-PGC-1α-derived MuSCs grown in proliferating conditions, while increased in transgenic cell-derived myotubes only, being consistent with the induction of *Mck* expression (Figure 2E).

Considering the accumulation of MuSCs in the skeletal muscle of transgenic animals, we tested if such abundance could result in improved muscle regeneration in middle-aged (12-months old) MCK-PGC-1α mice exposed to BaCl_2_ injury. Basic histological analyses revealed neither macroscopic morphological changes nor improved myofiber cross-sectional area (CSA) in 3-months old (young adult) transgenic animals compared to WT mice at 14 dpi (Figure 2F, 2G). Similarly, PGC-1α overexpression did not accelerate myofiber CSA recovery in middle-aged animals (Figure 2F, 2G). Overall, the metabolic switch imposed by PGC-1α overexpression impacts on MuSC number and increases their myogenic potential *ex vivo* but is unable to force regeneration *in vivo*.

### Transplantation of MCK-PGC-1α-derived cells recapitulates the oxidative phenotype in the skeletal muscle of WT mice

Since constitutive PGC-1α overexpression did not impinge on *in vivo* regeneration, whereas MuSCs from MCK-PGC-1α mice displayed enhanced *ex vivo* myogenic capacity (Figure 2C), the possibility that transplantation of myogenic progenitors derived from transgenic animals could contribute to muscle regeneration *in vivo* was investigated. To this purpose, mononucleated cells were isolated from hindlimb muscles of WT and MCK-PGC-1α donor animals and injected into the skeletal muscle (injured or intact) of WT recipients, according to the protocol described in Figure 3A. At 14 dpi, muscles transplanted with WT cells no longer showed morphological evidence of ongoing regeneration (Figure 3B) and presented with fully recovered CSA and a similar number of myofibers to non-injured muscles (Figure 3C, 3D). Consistently, no alterations on myogenin (MyoG) protein content (Figure 3E) or *Myh3* gene expression (Figure S3A) were observed. Nevertheless, Pax7 expression was increased in the injured group (Figure 3E), suggesting an increase in the MuSC pool. A completely different phenotype, with central myonuclei, cell infiltrate, reduced myofiber CSA and increased expression of Pax7 and MyoG was observed in both non-injured and injured muscles receiving MCK-PGC-1α cells (Figure 3B-C, 3E). Moreover, an increased number of myofibers was reported in injured muscles upon MCK-PGC-1α-derived cell injection (Figure 3D). This observation, associated with the parallel induction of *Myh3* expression (Figure S3A), was suggestive of new myofiber formation. Regarding the mitochondrial status, injured muscles transplanted with WT cells showed a complete recovery of oxidative/glycolytic myofiber distribution (Figure 3F) together with the persistent accumulation of PGC-1α and COX-IV proteins (Figure 3G). Strikingly, intact muscles injected with MCK-PGC-1α cells revealed a homogeneous SDH staining resembling the phenotype observed in transgenic animals (Figure 3F, S2B), suggesting that MCK-PGC-1α-derived myogenic progenitors might have fused into non-regenerating host myofibers, recapitulating the metabolic phenotype of transgenic mice. Additionally, a similar metabolic conversion was observed in muscles injured previous to transplantation with transgenic cells (Figure 3F). Consistent with these observations, PGC-1α and COX-IV protein levels increased in both uninjured and injured recipient muscles (Figure 3G).

**Figure 3.**
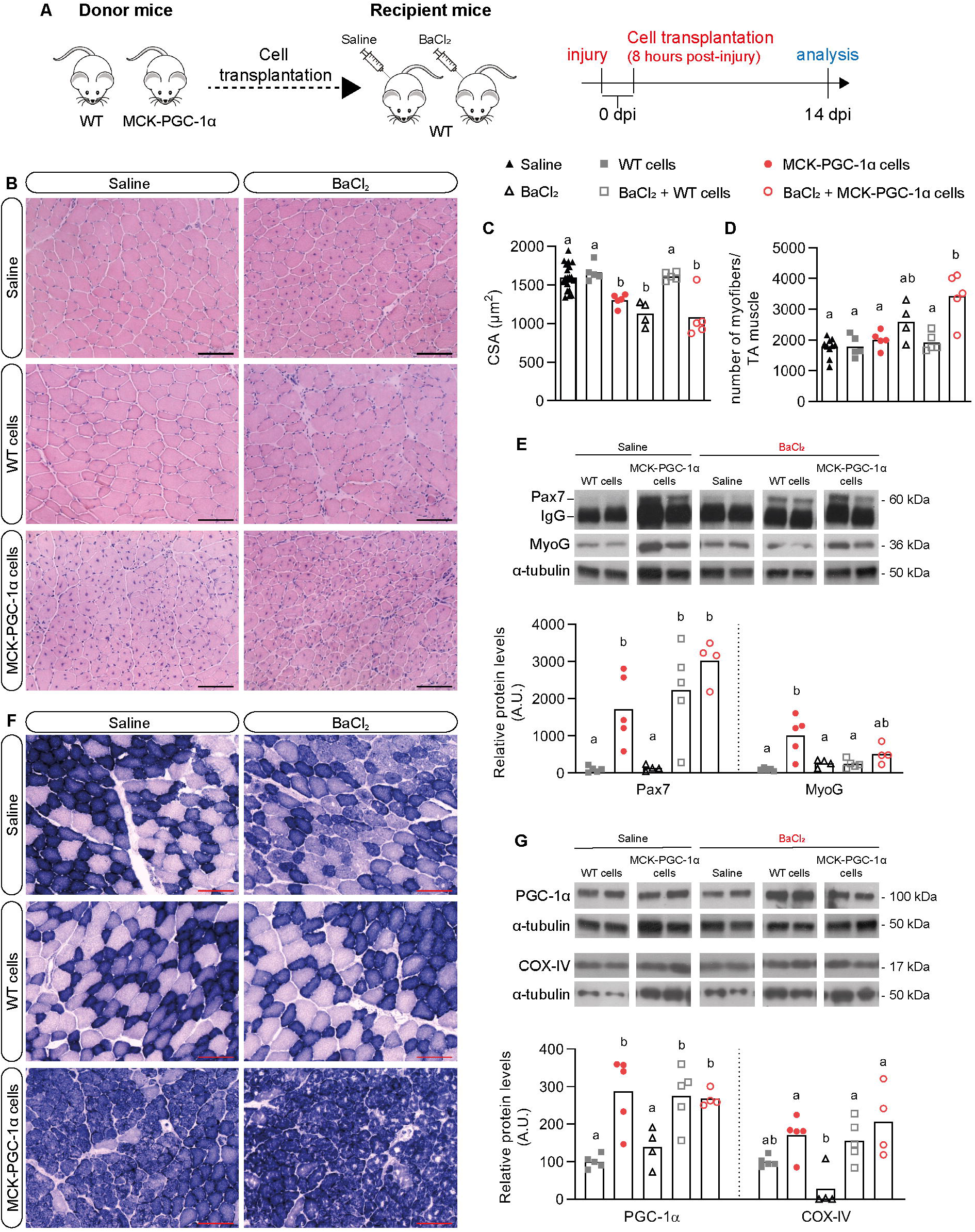
MCK-PGC-1α cell transplantation recapitulates the transgenic phenotype even in a non-regenerative environment. (A) Schematic representation of the transplantation experiment. Mononucleated cells were enzymatically isolated from hindlimb muscles of donor mice WT and MCK-PGC-1α mice and were transplanted to intact or injured TA muscles of recipient WT animals. Injured TA muscles were injected with 1.2% BaCl_2_ approximately 8 hours before cell transplantation. Groups of recipient mice consisted of n=5 animals, except for BaCl_2_ group with n=4 mice, and were sacrificed 14 dpi; (B) Representative images of H&E staining (scale bar: 100 μm); (C, D) CSA and total number of myofibers in a single TA muscle; (E, G) Representative western blotting bands and densitometry analysis of Pax7, MyoG, PGC-1α and COX-IV proteins. Total protein load was normalized by means of α-tubulin protein expression. Pax7, MyoG and PGC-1α share the same α-tubulin normalization images since they were analyzed on the same gel. Groups with distinct letters are statistically different (One-way ANOVA + Tukey’s test); (F) Representative images of SDH staining (scale bar: 100 μm).

### Overexpression of PGC-1α inhibits the adipogenic drift in the regenerating muscle

Muscle regeneration is not only dependent on MuSCs but also on ISCs. Deficient FAP function compromises MuSC activation, contributing to fibrosis and adipocyte infiltration during faulty muscle regeneration^6,39^. Recently, new subpopulations of adipogenic regulators with the capacity of modulating adipogenesis have been identified in mammalian fat depots^40,41^. Moreover, cell populations with similar regulatory capabilities are found in the skeletal muscle as part of specific ISC subtypes^42^. To better investigate the effects of muscle-specific PGC-1α overexpression on both myogenic and non-myogenic cells, we isolated mononucleated cells from WT and MCK-PGC-1α muscles and analyzed the ISC distribution by flow cytometry. In comparison to WT mice, transgenic animals presented increased Sca1^+^ cells associated with CD142 co-labeling, indicating an enrichment in adipogenesis regulators (Aregs; Figure 4A, B), a cell population described to exert anti-adipogenic functions^40,42^. On the contrary, the amounts of cells positive for CD55, a marker highly expressed in DPP4^+^ pre-adipocytes^41^, was lower in MCK-PGC-1α mice than in WT animals (Figure 4A, C).

**Figure 4.**
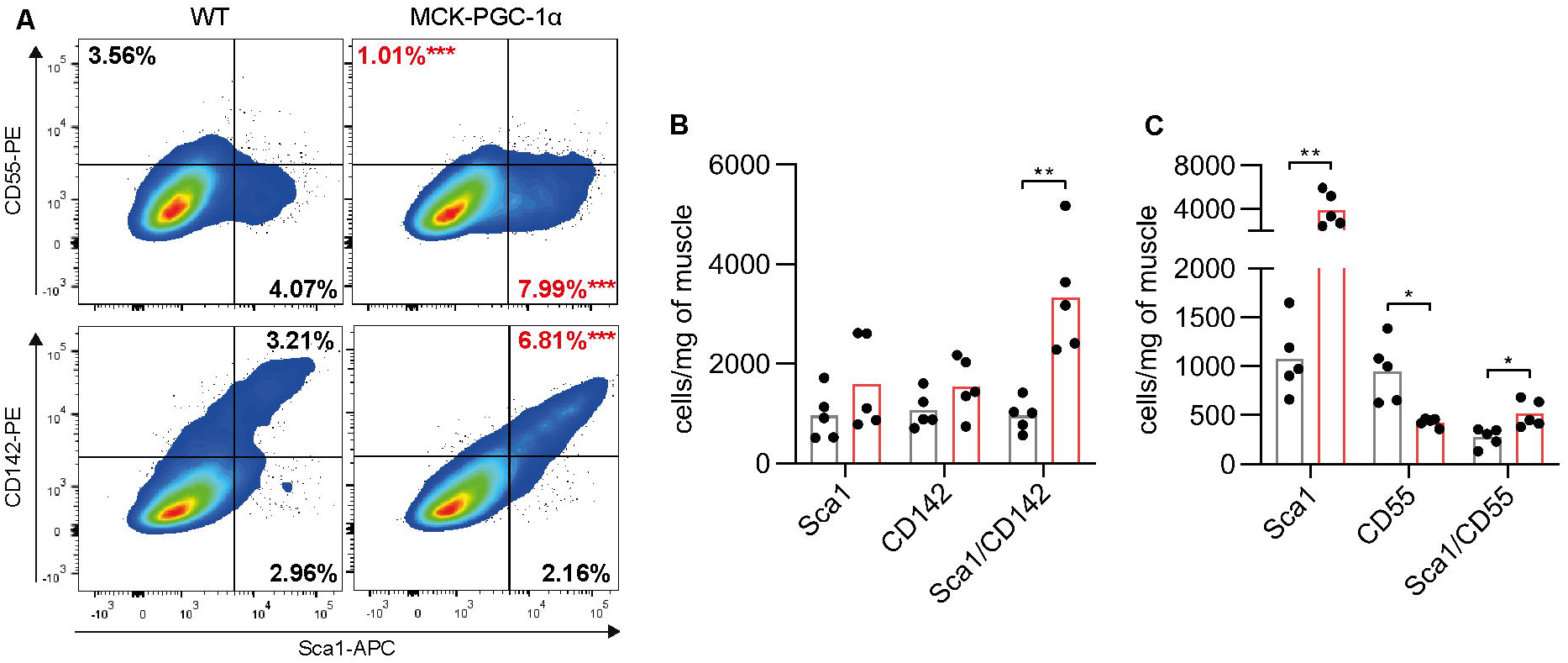
PGC-1α promotes the accumulation of Aregs against pre-adipocytes in the skeletal muscle. (A) Representative panels of gate distribution and relative quantification of positive events (cells) according to anti-Sca1, anti-CD55 or anti-CD142 antibody binding. Numbers in gates represent the mean percentage of cells by each specific labeling combination; (B) Sca1+, CD142+ and Sca1+/CD142+ cells normalized by total number of isolated cells by mg of tissue; (C) Sca1+, CD55+ and Sca1+/CD55+ cells normalized by total number of isolated cells by mg of tissue; Significance of the differences: *p<0.05, **p<0.01 *vs* WT group (Student’s t-test).

The altered proportions of Aregs and CD55^+^ cells suggested that the propensity to accumulate adipocytes after injury was altered in the skeletal muscle of PGC-1α overexpressing animals. To this purpose, 50% glycerol (Gly) was injected in the TA muscle of young WT and MCK-PGC-1α mice, reproducing a muscle injury model that promotes adipogenic infiltration^43^. As expected, a regenerative-like phenotype (Figure 5A) and the induction of *Myh3* expression was observed in Gly-injected TA muscles (Figure 5B). Injured muscle histology showed intramuscular white spots (Figure 5A), overlapping with Oil Red O (ORO) staining (Figure 5C). Additionally, RT-qPCR analysis revealed that adipocyte-related genes *Plin1*, *Adipoq* and *Fadq4* were induced in the muscle of both WT and MCK-PGC-1α mice (Figure 5B), and perilipin immunofluorescence staining demonstrated the accumulation of intramyofibrillar adipocytes (Figure 5D). Notably, expression of *Cd55* was significantly induced only in WT mice after Gly injury, whereas *Cd142* showed a strong tendency to increase in injured muscles of both WT and MCK-PGC-1α mice (Figure 5B). Consistent with the reported reduction in pre-adipocytes and increase in Aregs, adipocyte infiltration was partially prevented in MCK-PGC-1α animals as assessed by perilipin-immunofluorescence and densitometric analysis of the perilipin positive area (Figure 5D, E), demonstrating blunted adipogenic differentiation in the skeletal muscle of transgenic animals.

**Figure 5.**
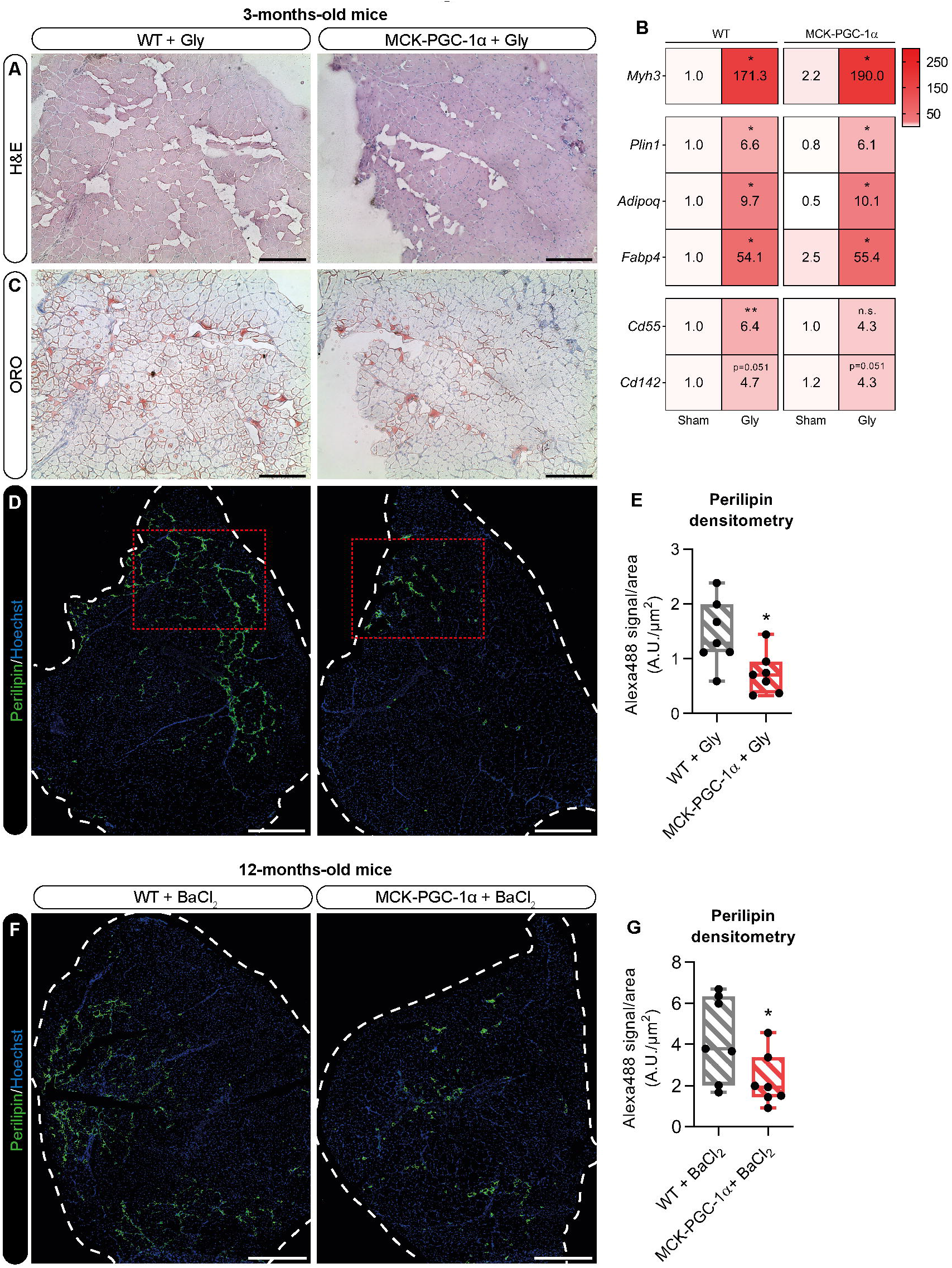
Muscle PGC-1α overexpression prevents the accumulation of intramuscular adipocytes. (A) Representative images of H&E staining (scale bar: 200 μm); (B) Heatmap showing relative expression of *Myh3*, *Plin4*, *Adipoq*, *Fapb4*, *CD55* and *Cd142* genes by RT-qPCR quantification. Data are normalized by *Actb* expression and displayed as relative expression (2^−ΔΔCt^; mean) *vs* WT-Sham group. Significance of the differences: *p<0.05 *vs* WT-Sham or MCK-PGC-1α-Sham group (Two-way ANOVA + Holm-Šídák test); (C) Representative images of ORO staining (scale bar: 200 μm); (D) Representative images of perilipin/Hoechst immunostaining (scale bar: 500 μm). Red-doted rectangles highlight the same area shown in panels C and D; (E) Densitometric quantification of Alexa488 signal, proportional to anti-perilipin antibody labeling. Significance of the differences: *p<0.05 *vs* WT + Gly (Student’s t-test); (F) Representative images of perilipin/Hoechst immunostaining of TA muscle of 12-months old WT (n=7) and MCK-PGC-1α mice (n=7) injected with BaCl_2_ at 14 dpi (scale bar: 500 μm); (G) Densitometric quantification of Alexa488 signal, proportional to anti-perilipin antibody labeling. Significance of the differences: *p<0.05 *vs* WT + BaCl_2_ (Student’s t-test).

To further investigate if muscle PGC-1α overexpression could reduce the adipogenic drift typically occurring during muscle regeneration in the elderly^44,45^, adipocyte infiltration in the skeletal muscle was also assessed in the above-mentioned 12-months old WT and MCK-PGC-1α animals undergoing BaCl_2_ injury (see data in Figures 2F, 2G). Perilipin-positive adipocytes were clearly detectable in injured muscles of middle-aged WT mice (Figure 5F). Consistent with the results obtained in young animals, older MCK-PGC-1α animals presented with a reduced accumulation of intramyofibrillar adipocytes in comparison to WT ones (Figure 5J). The present results demonstrate that the predominant Areg subpopulation found in PGC-1α overexpressing muscles is associated with a decreased propensity to adipocyte differentiation during defective regeneration in aged animals.

## DISCUSSION

Mounting evidence points to metabolism as a relevant factor in the regulation of stem cell quiescence, activation and differentiation in different tissues^46^. Specifically in the skeletal muscle, MuSC activation correlates with the induction of diverse metabolic pathways sustaining the high energy demand required for myoblast differentiation^13^, leading to an abrupt increase of mitochondrial activity and ATP availability^14^. In addition to supporting the increased energy demand, an increase in metabolic activity (both mitochondrial and non) generates metabolites that impinge on intracellular pathways that regulate stem cell function^47,48^. The current work provides evidence that an early, potentially PGC-1α-driven, activation of mitochondrial biogenesis in regenerating myofibers contributes to restore the metabolic phenotype of a healthy muscle. The main novelty is the qualitative and quantitative impact exerted by PGC-1α on the different muscle stem cell populations, including MuSCs and ISCs.

In the experimental conditions adopted in the current study, muscle regeneration in MCK-PGC-1α animals occurred with a similar kinetic in WT mice, despite the former show increased number of MuSCs which are endowed with enhanced *ex vivo* differentiation capacity. In this regard, PGC-1α effects on myogenesis *in vivo* could be amplified in conditions characterized by disrupted muscle regeneration such as muscular dystrophies, whereas in healthy conditions the availability of more MuSCs might not impact on an already efficient regeneration process. The other way around, syngeneic allograft of transgenic muscle-derived cells recapitulated the oxidative phenotype in both injured and intact recipient muscles. Considering that exogenous cell engraftment is rarely observed in the absence of host muscle damage^49^, the oxidative conversion observed in uninjured muscles supports the possibility that transgenic cells do have increased fusion capability and myogenic potential. The high MyoG expression in muscles transplanted with MCK-PGC-1α-derived cells could partially explain the increased fusion potential, consistently with recent evidence highlighting a central role played by MyoG in myoblast fusion^50^. However, the exact mechanism accounting for the improved ability of transgenic muscle progenitors to differentiate and fuse is far from being elucidated.

It is worth mentioning that the PGC-1α transgenic construct used for this study is under the control of the MCK promoter^51^, implying that only mature myofibers overexpress PGC-1α in intact skeletal muscle. However, MyoD was shown to be able to bind and activate the MCK promoter^52^. We have demonstrated that proliferating MuSCs obtained from transgenic animals do not present with overexpression of PGC-1α. However, we cannot rule out that the isolation procedure, involving MuSC removal from their niche and the consequent rapid induction of MyoD^53^, could result in a transient PGC-1α transgene induction critical to enhance progenitor cell engraftment. The promotion of mitochondrial respiration in myogenic progenitors was proven sufficient to endorse their differentiation potential^54^. In this line, an hypothetical early overexpression of PGC-1α in myogenic precursor cells could increase mitochondrial content in MuSCs, and would therefore partially recapitulate the increased differentiation capability observed by Haralampieva et *al*., in human myoblasts^55^. Nevertheless, our data support the idea that oxidative metabolism can be a potential tool to promote muscle repair when the regenerative capacity is impaired.

The present study also shows for the first time that myofiber PGC-1α overexpression prevents intramuscular fat accumulation. Intramuscular adipogenesis consequent to glycerol exposure is associated with an exacerbated inflammatory response as compared to the cardiotoxin-induced injury, with the activation of adipogenic regulatory networks and with reduction of fatty acid β-oxidation^56^. Data obtained in mice carrying a PDGFRα-reporter transgene indicate that most of the adipocytes that accumulate after glycerol injury arise from mesenchymal progenitors called FAPs^39^. However, whether FAPs can independently differentiate to adipocytes or require the interaction with other interstitial cells is still debated. The implementation of single cell transcriptomics to muscle regeneration has revealed that the Sca1^+^/PDGFRα^+^ cell pool includes a peculiar CD142^+^ population (Aregs) able to interfere with the propensity of CD142^-^ ISCs to undergo adipogenic differentiation^42^. The present study shows that the decreased accumulation of mature adipocytes occurring in the muscle of MCK-PGC-1α animals correlates with am unbalance between Aregs and CD55^+^ cells, the former being more abundant than the latter. These results are in line with previous observations demonstrating a regulatory role of Aregs on adipocyte differentiation and support the idea that the adipogenic fate of mesenchymal progenitors is modulated by the balance of specific ISCs populations that reside in the extracellular matrix^40,42^.

From a mechanistic point of view, it is conceivable that PGC-1α action on MuSCs and ISCs is driven by paracrine factors secreted by PGC-1α-overexpressing myofibers^33^. As an example, MCK-PGC-1α mice display high irisin levels. This latter is a polypeptide able to promote browning of the white adipose tissue^57^, thus being a factor potentially affecting adipocyte differentiation. Other hormonal mediators involved in modulating ISCs mitochondrial bioenergetics could also contribute to the anti-adipogenic drive reported in Gly-injured muscle of MCK-PGC-1α animals. For instance, impaired muscle regeneration and increased muscle adiposity induced by the genetic inhibition of α-Klotho mainly results from mitochondrial dysfunction in activated MuSCs^58^. This report could be consistent with the observation that in MCK-PGC-1α animals, characterized by a pro-oxidative muscle metabolic phenotype, adipogenic differentiation after glycerol injury or in BaCl_2_-treated middle-aged mice is inhibited.

The inhibition of adipogenesis in MCK-PGC-1α animals is particularly relevant to the study of diseases characterized by the progressive substitution of muscle mass with ectopic fat. Consistently, the results here reported reveal that adipocyte accumulation occurs in the injured muscle of middle-aged animals, and that such adipogenic drift is partially hindered in age-matched PGC-1α-overexpressing mice, entailing a promising connection between ISC dysfunction in aging and muscle oxidative metabolism. To fully validate these observations, further research should be performed on old and geriatric animals, as lipodystrophy occurring after regeneration increases with age and negatively affects muscle function^44,45^. Additionally, clarifying the mechanisms underlying the regulatory function exerted by PGC-1α on adipogenic populations is highly relevant to chronic muscle pathologies such as Duchenne muscular dystrophy or limb girdle muscular dystrophy 2B. Such diseases, indeed, are characterized by poor muscle morphology and increased disease severity that are associated with progressive replacement of muscle tissue with fat^59,60^.

In conclusion, the current results propose PGC-1α as a driver of muscle regeneration, achieved through the modulation of MuSCs and ISCs, favoring the myogenic lineage over the adipogenic one. Overall, the data here reported support the idea that harnessing muscle metabolism could become a therapeutic tool to treat muscle conditions characterized by impaired muscle regeneration.

## MATERIALS AND METHODS

All the reagents used in this study were obtained from Merck-MilliporeSigma (St. Louis, MO, USA) unless differently specified.

### Animals and experimental design

Experimental animals were cared for in compliance with the Italian Ministry of Health Guidelines and the Policy on Humane Care and Use of Laboratory Animals (NRC, 2011). The experimental protocols were approved by the Bioethical Committee of the University of Torino (Torino, Italy) and the Animal Welfare Committee of KU Leuven (Leuven, Belgium). Animals were maintained on a regular dark-light cycle of 12:12 hours with controlled temperature (18-23°C) and free access to food and water during the whole experimental period. Balb/c mice overexpressing PGC-1α in the skeletal muscle (MCK-PGC-1α) were generated by backcrossing C57BL/6-Tg(Ckm-Ppargc1a)31Brsp/j^51^ (The Jackson Laboratory, Bar Harbor, CA, USA) with WT Balb/c mice (Charles River, Wilmington, MA, USA). The resulting offspring was analyzed for the presence of the transgenic construct (forward: 5’-GCCGTGACCACTGCAACGA-3’ and reverse: 5’-CTGCATGGTTCTGAGTGCTAAG-3’) through Melt Curve Analysis (RT-qPCR) and selected for further crosses with WT Balb/c mice. The colony was maintained breeding mice as hemizygotes for 8 generations. During all intramuscular (i.m.) injections, the animals were anesthetized with 2% isoflurane in O2. The animals were sacrificed under anesthesia at specific time points. After intracardiac blood collection, euthanasia was applied by means of cervical dislocation. Skeletal muscles were excised, weighted, frozen in liquid nitrogen and stored at −80°C for further analyses.

The time course of muscle regeneration (animal experiment 1) was performed in WT 6-weeks old female mice receiving a local muscle injury (i.m. injection of 30 μl 1.2% BaCl_2_) in the TA muscle. Contralateral TA muscles were injected with filtered 0.9% NaCl solution and used as controls (Saline). Animals were euthanized at 7, 10, 13 and 49 dpi. Similarly, TA muscle injury of both WT and MCK-PGC-1α mice (animal experiment 2) was performed in 3-months and 12-months old female animals that were sacrificed at 14 dpi. Transplantation of isolated muscle-derived cells (animal experiment 3) was performed in WT 6-weeks old female mice previously injured in one of the TA muscles (approximately 8 hours before cell injection, see Figure 3A). Injected cells were isolated from the hindlimb muscles of either WT or MCK-PGC-1α male mice according to an adaptation of the protocol described by Costamagna et al.^11^(see below: *Whole muscle isolation*, *transplantation and flow cytometry analysis* section). Mice were euthanized at 14 dpi. As for adipogenesis study, glycerol injury (animal experiment 4) was performed by i.m. injection of 30 μl 50% glycerol sterile solution in the TA muscles of WT and MCK-PGC-1α 3-months old female mice. Animals were euthanized 21 dpi.

### Organ culture and myosin quantification

EDL and *Soleus* muscles from 6-weeks old male mice (WT and MCK-PGC-1α) were rapidly excised, rinsed in sterile PBS containing 5% antibiotics (9:1 penicillin/streptomycin:gentamicin) and digested in PBS containing 0.02% type I collagenase (C0130) for 1h at 37°C. Digested muscles were plated on matrigel-coated dishes in DMEM supplemented with 20% FBS, 10% horse serum, 0.5% chick embryo extract and 1% penicillin-streptomycin. Three days later, muscle remnant was removed and medium was replaced with proliferation medium (DMEM 4.5 g/l, 20% FBS, 10% horse serum, 1% chick embryo extract). After 5 days, the medium was replaced with differentiation medium (DMEM 4.5 g/l, 2% horse serum and 0.5% chick embryo extract). Primary myotube cultures were washed with PBS and fixed in acetone-methanol solution (1:1). Samples were then probed with anti-MyHC antibody (M4276) followed by labeling with secondary anti-mouse antibody Alexa Fluor 488 (A31627, Invitrogen, Carlsbad, CA, USA). Subsequently, myotubes were lysed with RIPA buffer (50 mmol/l Tris-HCl (pH 7.4), 150 mmol/l NaCl, 1% Nonidet P-40, 0.25% sodium deoxycholate, 1 mmol/l phenyl-methylsulfonyl fluoride), sonicated and centrifuged at 3000 rpm for 5 min. The pellet was discarded and fluorescence intensity of the supernatant was assessed (Alexa Fluor 488: ~495 nm excitation, ~519 nm emission; DAPI: ~358 nm excitation, ~461 nm emission).

### Whole muscle cell isolation, transplantation and flow cytometric analysis

Hindlimb muscles from 6-weeks old male mice (WT and MCK-PGC-1α) were mechanically minced using a scalpel, washed with 5% antibiotics and enzymatically digested with 0.02% type I collagenase (C0130) and 0.06% pancreatine (P3292) upon shaking for 1 hour at 37°C. The suspension was then filtered using 70 μm strainers (Falcon), the digestion was blocked with fetal bovine serum and cell suspension was kept at 37°C while the solid part was re-digested for additional 30 minutes repeating the steps above. For cell transplantation, the cell suspension was centrifuged for 5 minutes at 400 *g* at room temperature, resuspended in sterile saline solution and the equivalent amount of cells obtained from one TA donor muscle was injected in either injured or intact TA muscle of WT recipient mice. As for flow cytometric analysis, hindlimb muscles were enzymatically digested with 0.1% type II collagenase (C6885). Isolated cells were resuspended in growing medium (DMEM 4.5g/l glucose + 10% FBS) and kept at 37°C. After 4 hours, samples were probed with different combinations of conjugated primary antibodies (Table S1) and analyzed by flow cytometry (Canto II AIG, BD Biosciences, Franklin Lakes, NJ, USA). Finally, cultured MuSCs were isolated using two sequential enzymatic digestions while shaking at 37°C: a first incubation with 0.04% collagenase II (C6885) for 45 minutes, followed by 30 minutes digestion with 0.1% collagenase/dispase (11097113001, Roche Diagnostics, Mannheim, Germany). MuSCs were selected using the Satellite Cell Isolation Kit mouse (130-104-268, Miltenyi Biotec, Bergisch Gladbach, Germany), seeded at 2000 cells/cm^2^ and grown for 3 days in growth medium (GM) containing 20% horse serum, 3% chick embryo extract, 1% HEPES, 1% glutamine and 1% penicillin-streptomycin. To induce myotube formation, GM was replaced with differentiating medium (DM) containing 2% horse serum and kept for 3 days.

### Muscle histology

TA muscles were frozen in melting isopentane cooled in liquid nitrogen and stored at −80°C. Transverse sections of 10 μm from the midbelly region were cut on a cryostat, left at room temperature for 10 minutes and stored at −80°C for later staining. Hematoxylin/eosin staining (H&E) was performed following standard procedures. SDH staining was performed by incubating pre-warmed sections with SDH reagent (1 mg/ml NTB, 27 mg/ml sodium succinate in PBS) at 37°C for 20 minutes. Slides were then rinsed twice with PBS, dehydrated using ethanol scale and xylene, mounted using Eukitt Quick-hardening mounting medium (03959). ORO staining was performed on sections fixed with 4% paraformaldehyde by incubating Oil Red O solution (0.5% in propylene glycol) at 60°C for 15 minutes. After washing with 85% propylene glycol for 5 minutes, slides were counterstained with hematoxylin for 1 minute, air-dried and mounted with glycerol-PBS (3:1). Images were captured by a Leica DM750 optical microscope (Leica Camera AG, Wetzlar, Germany).

For immunofluorescence, the sections were fixed in 4% paraformaldehyde for 15 minutes, rinsed in PBS and probed with primary anti-laminin (1:100, L9393) or anti-perilipin A/B (1:200, P1873) antibodies. Detection was performed using Alexa Fluor 488-conjugated secondary antibody (Invitrogen) and nuclei were counterstained with Hoechst 33342. Slides were mounted with glycerol-PBS (3:1) and fluorescence images were captured by an Axiovert 35 fluorescence microscope (Zeiss, Oberkochen, Germany) without altering light exposure parameters. Whole TA muscle sections were generated using GimpShop software. Perilipin densitometry on whole muscle sections was quantified using the ImageJ software.

### Isolation, retro-transcription and RT-qPCR quantification of mRNA

TA muscles were lysed in 1 ml of TRI Reagent and processed using the standard phenol-chloroform method. Briefly, 50 μm thick muscle sections were shaken in 1 ml of TRI Reagent for 30 minutes at 4°C, added 200 μl chloroform, mixed vigorously and centrifuged at 12000 *g* for 15 minutes. The RNA in the aqueous part was precipitated by 2-propanol and 70% ethanol, dried at room temperature and resuspended in sterile water. RNA concentration was quantified using Ribogreen reagent (Invitrogen). Total RNA was retro-transcribed using cDNA synthesis kit (Bio-Rad, Hercules, CA, USA) and transcript levels were determined by RT-qPCR using the SsoAdvanced SYBR Green Supermix and the CFX Connect Real-Time PCR Detection System (Bio-Rad). Every RT-qPCR was validated by analyzing the respective melting curve. Primer sequences are given in Table S2. Gene expression was normalized to *Actb* and results are expressed as relative expression (2^−ΔΔCt^).

### Western blotting

TA muscles were mechanically homogenized in RIPA buffer (PBS, 1% Igepal CA-630, 0.1% SDS) containing protease inhibitors (0.5 mM PMSF, 0.5 mM DTT, 2 μg/ml leupeptin, 2 μg/ml aprotinin), sonicated for 10 seconds at low intensity, centrifuged at 15000 *g* for 5 minutes at 4°C and the supernatant was collected. Total protein concentration was quantified with Bradford reagent (Bio-Rad), using BSA as protein concentration standard. Equal amounts of protein (15-30 μg) were heat-denatured in sample-loading buffer (50 mM Tris-HCl, pH 6.8, 100 mM DTT, 2% SDS, 0.1% bromophenol blue, 10% glycerol), resolved by SDS-PAGE and transferred to nitrocellulose membranes (Bio-Rad). The filters were blocked with Tris-buffered saline (TBS) containing 0.05% Tween and 5% non-fat dry milk and then incubated overnight with antibodies directed against specific proteins (Table S2). Peroxidase conjugated IgGs (Bio-Rad) were used as secondary antibodies. Quantification of the bands was performed by densitometric analysis using a specific software (TotalLab, NonLinear Dynamics, Newcastle upon Tyne, UK).

### Statistics

Data are presented using bar and dot plots (mean) or box and whisker plots (line: median, whiskers: min to max), unless differently stated. Data representation and evaluation of statistical significance was performed with Prism (version 7, GraphPad) software. Normal distribution was evaluated by Shapiro-Wilk test. The significance of the differences was evaluated by appropriate two-sided statistical tests, being Student’s “t”-test or analysis of variance (ANOVA) for normal distribution, and Mann–Whitney test or Kruskal–Wallis test for non-normal distribution. ANOVA and Kruskal–Wallis tests were followed by suitable post hoc analysis.

## ACKNOLODGEMENTS

This work was supported by World Anti-doping Agency (WADA; Grant IOC15E01PC, P.I. P. Costelli), Fondazione CARIPLO (Grant 2017-0604 to F. Penna), Fondazione AIRC (IG 2018—ID. 21963 project, P.I. F. Penna), University of Torino (ex-60% funds), Consorzio Interuniversitario di Biotecnologie (Contributi mobilità 2018 to M. Beltrà), Association française contre les myopathies (AFM)-Téléthon (European Post-doctoral fellowship n°20673; to D. Costamagna) and Fonds Wetenschappelijk Onderzoek (FWO; Grant G0D4517N to M. Sampaolesi).

## AUTHOR CONTRIBUTIONS

MB designed, performed and analyzed the data of the majority of the experiments under the guidance of MS, F Penna and PC; MB, F Penna and PC wrote and edited the manuscript, whereas D Costamagna, RD, VM, D Coletti and MS revised the manuscript; F Pin conducted the experiments regarding *ex vivo* tissue culture and BaCl_2_ injection in young animals shown in Figure 2; D Costamagna and RD contributed to the isolation and flow cytometry analysis of cells isolated from hindlimb muscles shown in Figure 2D and Figure 4; AR contributed to the isolation, expansion and differentiation of MuSCs analyzed in Figure 2E; RB, LGC and AI contributed to cell isolation and transplantation, as well as conducted some of the downstream procedures related to Figure 3.

