## Supplementary figures and images for "PGC-1α in the myofibers regulates the balance between myogenic and adipogenic progenitors affecting muscle regeneration"

### Figure S1, S2, S3

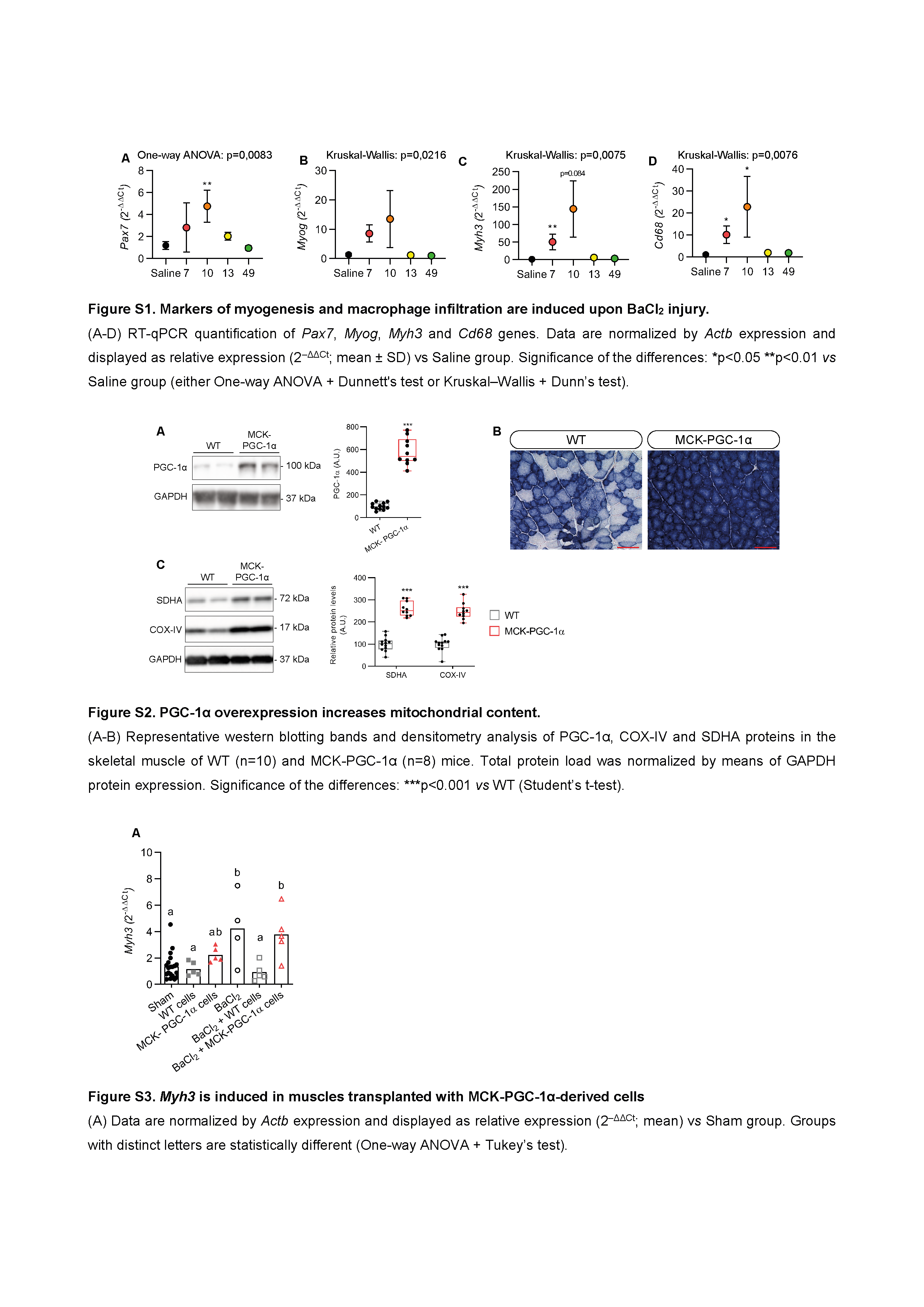
